# Cell-scale porosity in microporous annealed particle (MAP) scaffolds modulates immune response and promotes formation of innervated muscle fibers in volumetric muscle loss injuries

**DOI:** 10.1101/2024.05.31.596879

**Authors:** Areli Rodriguez Ayala, George Christ, Donald Griffin

## Abstract

Volumetric muscle loss (VML) is caused by severe traumatic injuries to skeletal muscle and is characterized by the irreversible loss of contractile tissue and permanent functional deficits. VML injuries cannot be healed by endogenous mechanisms and are exceptionally difficult to treat in the clinic due to the excessive upregulation of the inflammatory response, which leads to fibrosis, denervation of muscle fibers, and impaired regeneration. These injuries lead to long-term disability. Using a rodent model of VML in the tibialis anterior, this study presents microporous annealed particle (MAP) hydrogel scaffolds as a biomaterial platform for improved muscle regeneration in VML injuries, specifically highlighting the benefits of cell-scale porosity. In contrast to bulk (i.e., nanoporous) hydrogel scaffolds, MAP scaffolds promote integration by avoiding the foreign body response, decreasing the rate of implant degradation, and shifting macrophage polarization to favor regeneration. In addition, cell migration and angiogenesis throughout the implant precede the degradation of MAP scaffolds, including the formation of muscle fibers and neuromuscular junctions within MAP scaffolds prior to degradation. These fibers and junctions continue to develop as the implant degrades, indicating that MAP hydrogel scaffolds are a promising therapeutic approach for VML injuries.

## 1. Introduction

Volumetric muscle loss (VML) is defined pathologically as the irreversible loss of muscle tissue and function following severe injury to skeletal muscle. The scale and severity of VML-associated injuries result in chronic dysregulated inflammation and fibrosis that surpass the endogenous regenerative capacity of skeletal muscle. The exacerbated fibrosis within the muscle disrupts the functional and structural properties of the remaining tissue and can increase the susceptibility of surrounding tissue to further injury^1, 2^. Treatment strategies that are currently available in the clinic, such as autologous tissue transfer (i.e., muscle grafts) and physical therapy, are not effective at restoring muscle function. Additionally, autologous muscle grafts are invasive, may cause donor site morbidity, can have complications that result in complete graft failure (e.g., necrosis and infections), and are not a viable option for all patients^3–6^. Physical therapy alone does not provide significant improvements in function, either^7, 8^. Consequently, VML patients experience long-term, progressive disability. Along with the loss of contractile tissue, VML injuries destroy supporting structures that are essential to muscle function and regeneration (e.g., basal lamina, neuromuscular junctions, nerves, satellite cells, and blood vessels) in the area of the defect, further complicating recovery; therefore, an effective treatment must restore not only the lost muscle fibers, but all supporting structures as well.

A wide variety of biomaterial-based approaches have been used to support VML recovery, including decellularized matrices^8–10^, electrospun scaffolds^11^, bioactive glass^12^, nanomaterials^11, 13–18^, and hydrogels^19–26^. While biomaterial-assisted therapies are promising tools to provide a guided healing environment for the restoration of lost tissue and function in VML, they have yet to achieve clinical significance.^27^ In order to effectively promote regeneration, a biomaterial implant must overcome the natural predisposition to fibrosis by addressing the severe, local immune dysregulation; provide a scaffold to rebuild functional support structures; and promote the regeneration of organized, contractile muscle fibers. Of the biomaterial classes listed above, hydrogels are one of the most commonly used for VML due to their chemical and physical versatility, which allow them to match skeletal tissue properties. Hydrogels for muscle regeneration have been made from natural and synthetic polymers, and have been applied in various forms (e.g., bioprinting^19–21^, electrospinning^22^, sponges^23–25^, and granular hydrogels^26^). Like other biomaterials, hydrogels face an additional obstacle in the form of a foreign body reaction (FBR), which is a form of chronic inflammation that occurs at the interface of the implant and ultimately causes its fibrotic encapsulation. Therefore, biomaterial-based scaffolds must mitigate the FBR to integrate with host tissue and benefit VML regeneration. Outside of VML treatment, general biomaterial strategies to manage FBR include altering the material stiffness^28^, modifying the material chemistry^29–31^, attaching bioactive molecules^32–34^, and releasing anti-fibrotic drugs from the implant^35–39^. Another interesting strategy is to incorporate cell-scale porosity, which has been to shown to exhibit immunomodulatory properties, including a decrease in FBR^40, 41^.

Along with reduced inflammation, the regeneration of neuromuscular junctions (NMJs) and motor neurons within the injury site is essential to full functional recovery (rather than functional fibrosis^42, 43^) in a muscle that has sustained a VML injury. NMJs are synapses between motor neurons and skeletal muscle, and are required for normal neuromuscular function *in vivo*.^44^ Disruptions to NMJ structure and stability, such as denervation (i.e., loss of the presynaptic terminal), in multiple pathological conditions (e.g., neurological disorders, nerve injury, VML, and chronic disuse) lead to muscle atrophy and loss of function.^44–48^ Conversely, the extent of re-establishment of NMJs after peripheral nerve injury has a strong positive correlation with functional recovery.^46^ Reinnervation is of particular importance in VML due to the secondary denervation of muscle fibers that follows the primary loss of nerves and NMJs at the time of injury, which results in significant denervation of NMJs by 48 days post-injury and further functional decline.^47^ Therefore, an effective therapy for VML must promote proper NMJ formation and reinnervation of muscle fibers within the defect area. However, few studies of biomaterial-based therapies for VML have quantified the re-establishment of NMJs within or in the vicinity of the defect. The small number of studies that have documented regenerated NMJs in and around the injury have included cells in the scaffold^22, 49, 50^, have implanted tissue engineered constructs^51, 52^, or have added an rehabilitative component (i.e., exercise)^53^. To date, we have not found any examples in the literature of acellular scaffolds that significantly increased the number of NMJs within VML defects in sedentary animals.

Here, we explore the treatment of VML injuries with Microporous Annealed Particle (MAP) scaffolds, which are injectable hydrogel constructs that start as a slurry of spherical microgels that become bonded together (i.e., annealed) *in situ* to form a structurally stable scaffold with cell-scale porosity. This biomaterial platform has been used for multiple regenerative engineering applications, including dermal wound healing^54, 55^, brain recovery from stroke^56, 57^, cardiac muscle recovery following myocardial infarction^58^, vocal fold augmentation^59, 60^, acute cartilage defect healing^61^, recovery from spinal cord injury^62^, and delivery of mesenchymal stem cells^63^. Additionally, previous work has shown that MAP scaffold porosity has immunomodulatory properties, as demonstrated by their lack of fibrous encapsulation^60, 64^, mitigation of glial scarring^57, 62^, and decreased presence of immune cells with inflammatory phenotypes^54, 57, 62, 65^. We tested the potential of acellular MAP scaffolds to address many of the challenges of regeneration after a VML injury in a rodent model, evaluating the effect of microporosity specifically by comparing MAP scaffolds to nanoporous bulk scaffolds with the same chemical composition and stiffness. In addition, the hydrogel formulation we have used in this study has negligible bioactivity of its own, with only a common cell adhesion peptide motif (RGD) included. This study is the first to use MAP scaffolds for skeletal muscle regeneration, and we provide evidence that the inclusion of microporosity in a biomaterial-based treatment for VML can modulate the local immune response, mitigate the FBR and prevent encapsulation of the implant, promote myofiber formation within the scaffold, alter material degradation rates, and support the formation of NMJs in the defect area.

## 2. Results and Discussion

### 2.1. Synthesis and Characterization of MAP hydrogel

The nanoporous and MAP scaffolds used in this study were formulated to be chemically identical following application, so as to isolate the effects of porosity and geometry on tissue-biomaterial interactions. The precursor solution consisted of a 3.0 wt% polyethylene glycol (PEG)-maleimide backbone crosslinked with a matrix metalloprotease (MMP)-cleavable peptide linker; RGD peptide to promote cell adhesion; a custom-made, heterofunctional 4-arm PEG-maleimide/methacrylamide macromer^66^ to accelerate photo-annealing (Fig. **1a**); and a biotin-maleimide label to allow for downstream fluorescent identification. The precursor solution was used without further processing for the nanoporous implants, while MAP microgels were generated using a step-emulsification microfluidic device as previously described (Fig. **1b**).^67^ This particular method of microgel production allows for tighter control of microgel size and the generation of a monodisperse population of particles. Variables such as the channel height of the microfluidic device, the flow rate of the precursor solution, and the flow rate of the oil phase can be modulated to obtain a target size and variability. In addition, the composition of the precursor solution was tailored to fit the biomechanical properties of myofibers by adjusting the PEG content of the network to match compressive stiffnesses. Upon gelation, the 3.0% PEG (w/v) precursor solution approximated the optimal stiffness of a substrate for myotube differentiation (12 kPa^68^), with a Young’s modulus of 11.35 ± 1.18 kPa (Fig. **1c**). Further, the channel height of the microfluidic device, precursor solution flow rate, and oil flow rate used for microgel generation (11.4 μm, 3.75 ml hr^-1^, and 7.5 ml hr^-1^, respectively) were chosen to create a monodisperse population of microgels with a particle diameter of 86.26 ± 14.97 μm and a polydispersity index (PDI) of 1.03 (Fig. **1d**).

**Figure 1.**
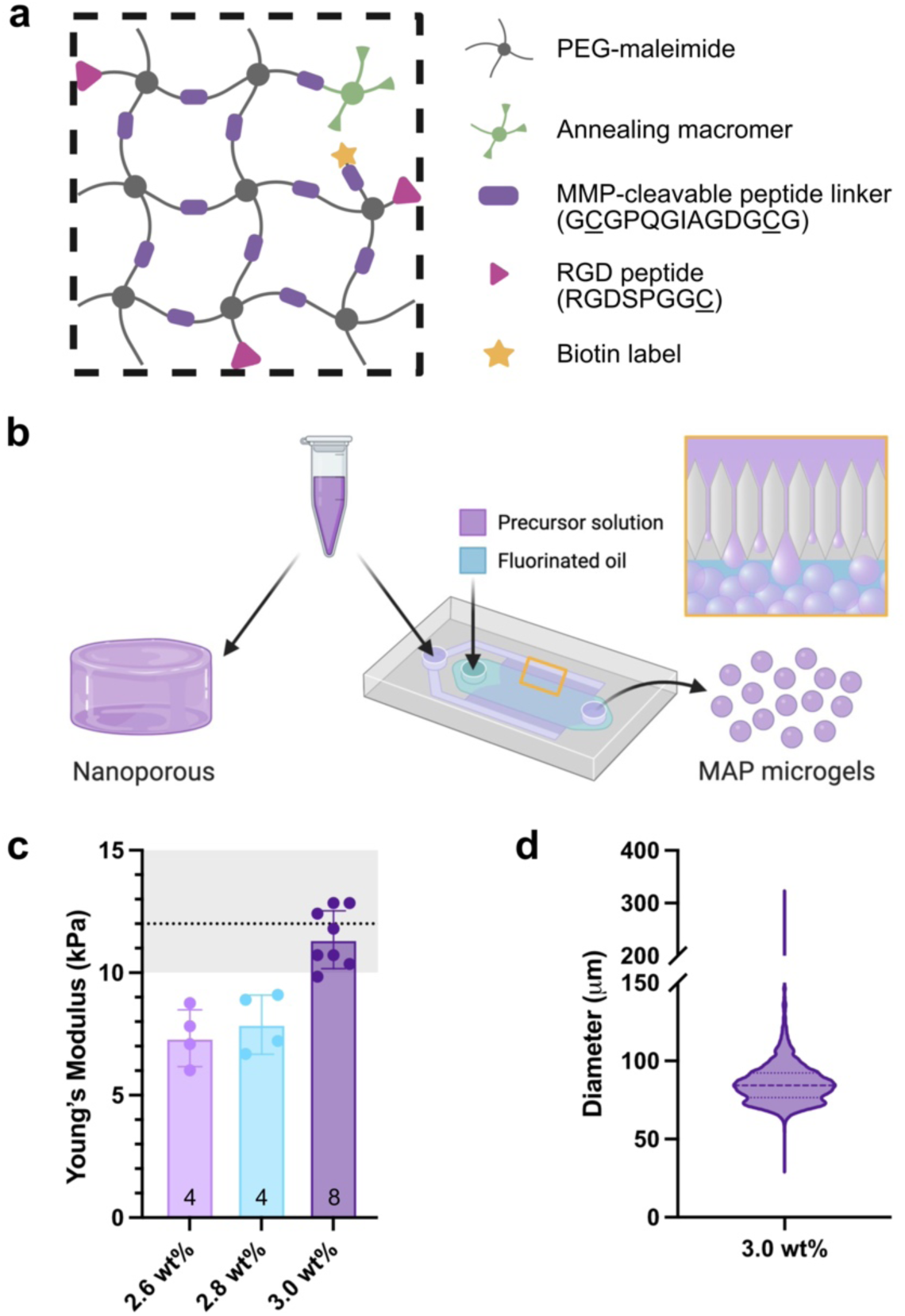
Hydrogel composition and microgel generation. (a) The hydrogel formulation contained PEG-maleimide, an enzymatically degradable peptide crosslinker, a cell adhesion peptide motif, and an annealing macromer. (b) The same precursor solution was utilized for nanoporous implants and MAP microgels, which were generated by step-emulsification in a microfluidic device. (c) The PEG content of the precursor solution was determined by compressive stiffness post-gelation, with a target of 12 kPa. (d) Microgels used to create MAP scaffolds were monodisperse (particle diameter of 86.26 ± 14.97 μm and a PDI of 1.03).

### 2.2. Tibialis anterior VML defect creation and treatment

Animals were weighed and function-tested before undergoing any surgical procedures, and the maximum torque produced by each animal upon stimulation of the peroneal nerve was normalized by body weight. The three treatment groups for this study were MAP hydrogel, nanoporous hydrogel, and an untreated negative control. To characterize the regeneration of muscle at different stages, this study had two cohorts with endpoints at 4 weeks (n=8 per group) and 12 weeks (n=7 per group) post-injury, respectively (Fig. **2a**). The first endpoint was chosen to coincide with the tail-end of the remodeling phase of an acute muscle injury.^69^ The second endpoint was chosen to evaluate late-stage regeneration and more accurately assess functional recovery.^70^ After calculating their baseline function relative to body weight, animals were assigned to treatment groups that maintained statistically equal means for normalized function across groups (MAP: 117.85 ± 9.28 N mm kg^-1^, Nanoporous: 118.58 ± 7.20 N mm kg^-1^, Untreated: 116.51 ± 10.68 N mm kg^-1^) (Fig. **2b**). Immediately prior to surgery, the theoretical mass of the tibialis anterior (TA) was calculated for each animal following a previously published formula^71^, and this value was used to create a defect of approximately 20% (w/w) in the middle section of the TA. Defects were consistent across groups (MAP: 21.53 ± 0.71%, Nanoporous: 21.46 ± 0.82%, Untreated: 21.47 ± 0.65%) (Fig. **2c**). After creation of the injury, hydrogel treatment or saline were applied to the defect with a pipette (Fig. **2d**). Nanoporous scaffolds were allowed to gel *in situ* while MAP microgels were annealed *in situ* to form a bulk scaffold, followed by closing of the fascia and skin with sutures.

**Figure 2.**
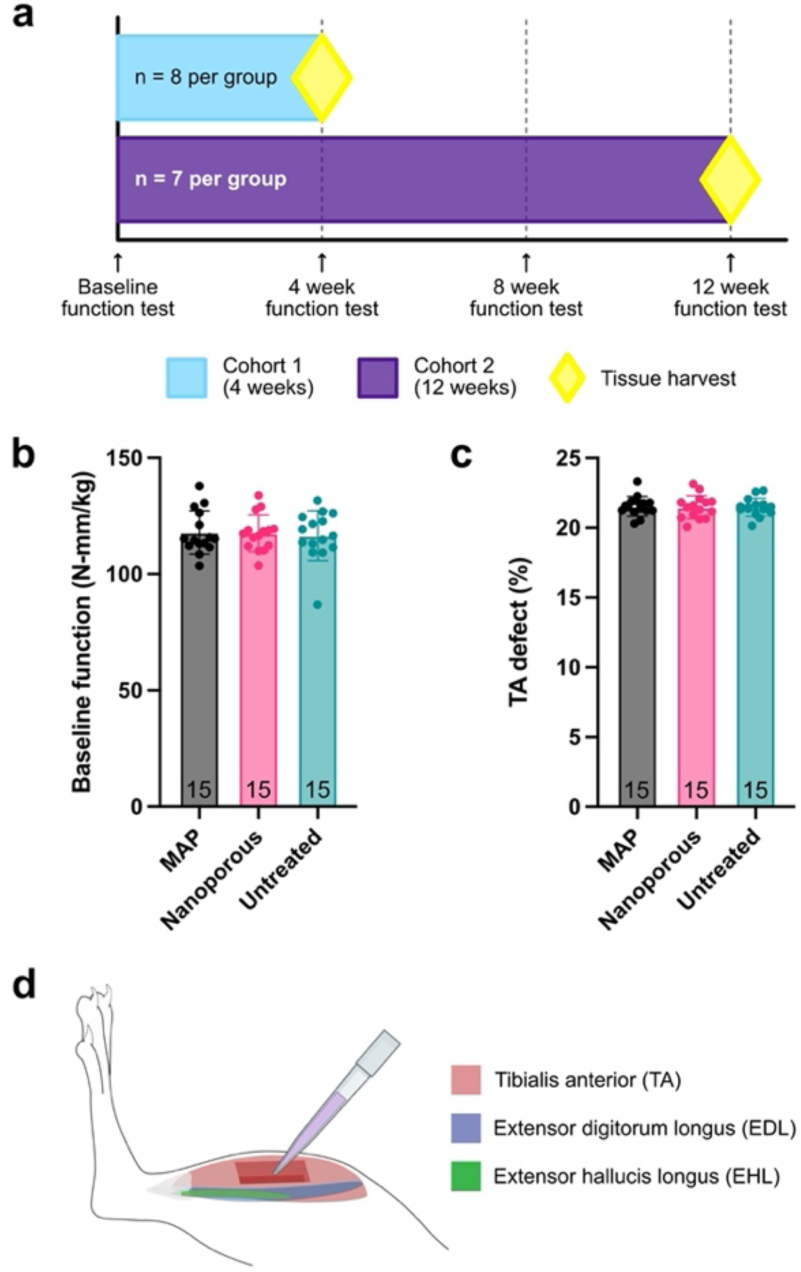
Study design and defect creation. (a) The study consisted of two cohorts to study mid-stage and late-stage muscle regeneration (b) Animals were assigned to treatment groups such that baseline normalized function was consistent across groups, with no significant differences. (c) Approximately 20% of the TA was surgically removed and the size of the defect was consistent across treatment groups, with no significant differences. (d) Synergist muscles (EDL and EHL) were removed, followed by creation of the defect in the middle third of the TA. Treatments were applied to the defect with a pipette. Statistics: One-way ANOVA, multiple comparisons post-hoc tests (Tukey HSD)

### 2.3. Microporosity counterintuitively decreases rate of implant degradation

Magnetic resonance imaging (MRI) was used to quantify the remaining scaffold in the defect and thus determine the effect of porosity on scaffold degradation. The animals were imaged at 4 weeks and 12 weeks post-injury. Earlier time points were not used for this analysis due to the prevalence of edema at the site of VML injuries at early stages. Due to their high water content, hydrogel implants produce bright signal in T2-weighted images. To track our implants, cross-sections of the lower leg were obtained such that the tibialis anterior and defect area could be easily identified using the tibia and fibula as landmarks (Fig. **3a**). Representative MR images (Fig. **3b**) show that both MAP and nanoporous implants (outlined) are clearly visible. In addition, persistent edema in the untreated defect (brighter regions in and around the tibialis anterior) is visible at 4 weeks and has resolved by 12 weeks, while there appears to be some edema remaining at both timepoints for the hydrogel groups. Interestingly, nanoporous implants had an increased degradation rate compared to MAP implants, with a significant difference in remaining scaffold volume at 4 weeks (Fig. **3c**). Although the difference in scaffold volume at 12 weeks is not significant, the trend is preserved. We believe this is due to the activity of local immune cells, suggesting that they are more active in response to nanoporous implants compared to MAP implants. Further, it is important to note that the hydrogel network does not occupy the entirety of a MAP scaffold due to the void spaces created by its microporosity. Approximately 64-74% (assuming random close packing of spheres or perfect packing of spheres, respectively) of the scaffold volume is material that can be degraded, while the rest is filled with fluid or tissue. Therefore, the difference in degradation rates of the hydrogel network is underestimated by overall volume measurements and would require the collection of additional data to be evaluated more accurately. This finding is noteworthy and could influence the design of future biomaterial-based therapies for VML, given that the effect of bioactive components introduced through implants is often limited by the permanence of the scaffold. Adding microporosity to an implant could be used to prolong the therapeutic window of bioactive components (e.g., growth factors, cytokines, pharmacologic agents, and extracellular vesicles) that are immobilized or encapsulated in a hydrogel and released upon degradation of the network. In some cases, it could potentially decrease the cost of including these bioactive components in the scaffold; a smaller quantity embedded in a microporous implant might be able to achieve the same effect as a larger quantity in a nanoporous implant.

**Figure 3.**
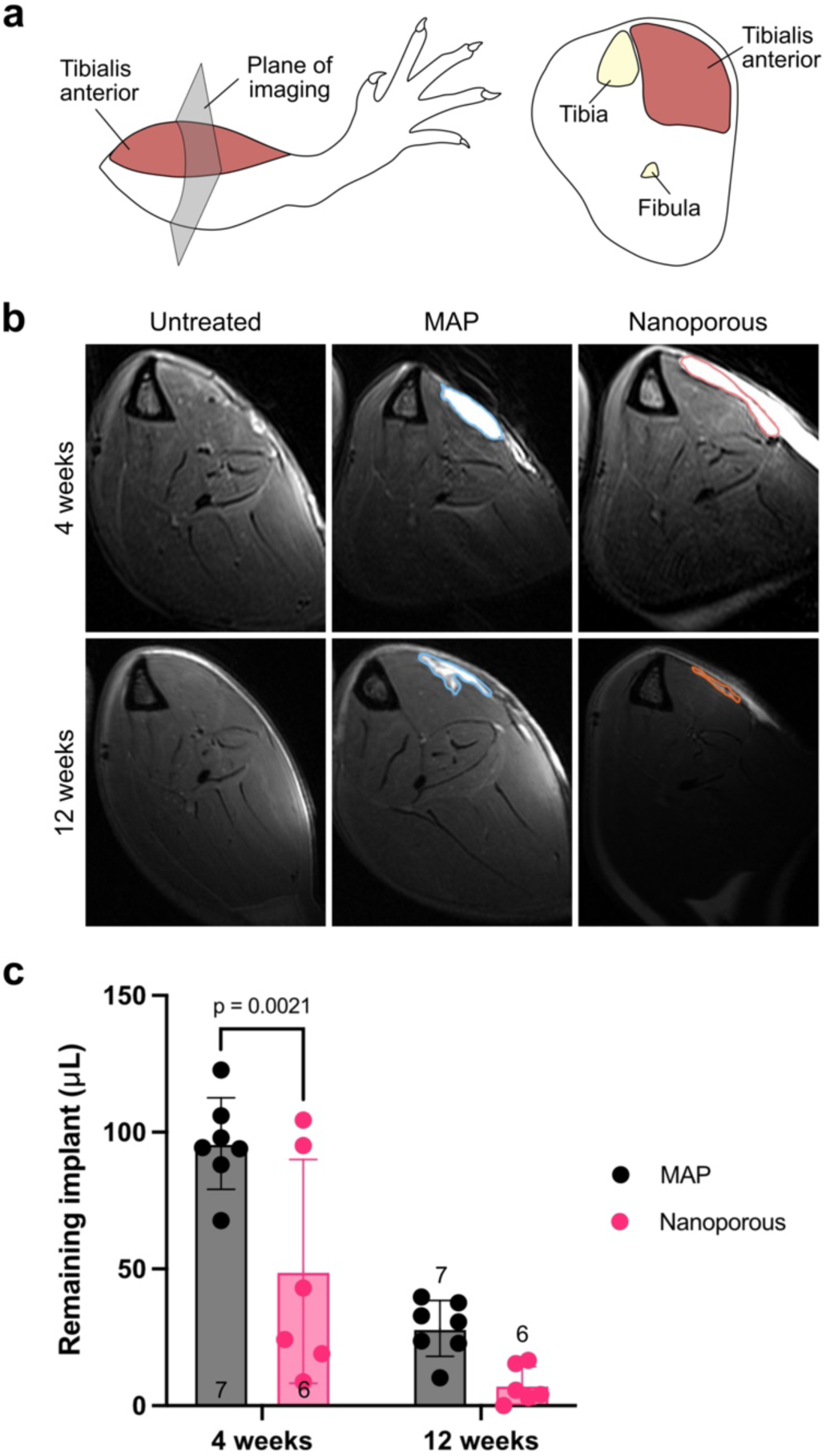
Quantification of implant retention by MRI. (a) Cross-sections of the lower hindlimb were imaged. The tibia, fibula, and TA were used as landmarks to locate the implant. (b) Representative images (single slice) of each treatment group at different timepoints. Areas of bright signal indicate high water content. Hydrogel implants (outlined) are localized to the TA. Edema may be found in areas surrounding VML injuries. (c) Volumetric quantification of the remaining implants, showing accelerated degradation of the nanoporous scaffolds. Statistics: Two-way ANOVA, multiple comparisons post-hoc tests (Šídák)

### 2.4. Microporosity lowers local inflammatory response to scaffold implant

Another important distinction in tissue-implant interactions between the two hydrogel treatments is related to the foreign body response. Picrosirius red staining was used to distinguish collagenous tissue (in red) from muscle fibers (in yellow). We observed the formation of fibrotic capsules around the nanoporous implants at both 4 weeks and 12 weeks post-injury, while no such capsule formed at either timepoint for MAP implants (Fig. **4a**). Additionally, newly formed fibers are visible within MAP scaffolds as early as 4 weeks, another strong indication that cellular migration and tissue integration has not been hindered by a FBR. Quantification of the capsule thickness (Fig. **4c**) showed that the capsule thickness was dynamic between 4 weeks and 12 weeks, exhibiting a significant difference in thickness with time (221.74 ± 119.58 μm at 4 weeks, 58.82 ± 63.73 μm at 12 weeks). The binary contrast in implant encapsulation between the two hydrogel treatments shows that the FBR is rendered negligible by the microporosity of MAP scaffolds, and therefore FBR avoidance does not require the inclusion of anti-fibrotic molecules in the implant’s composition.

**Figure 4.**
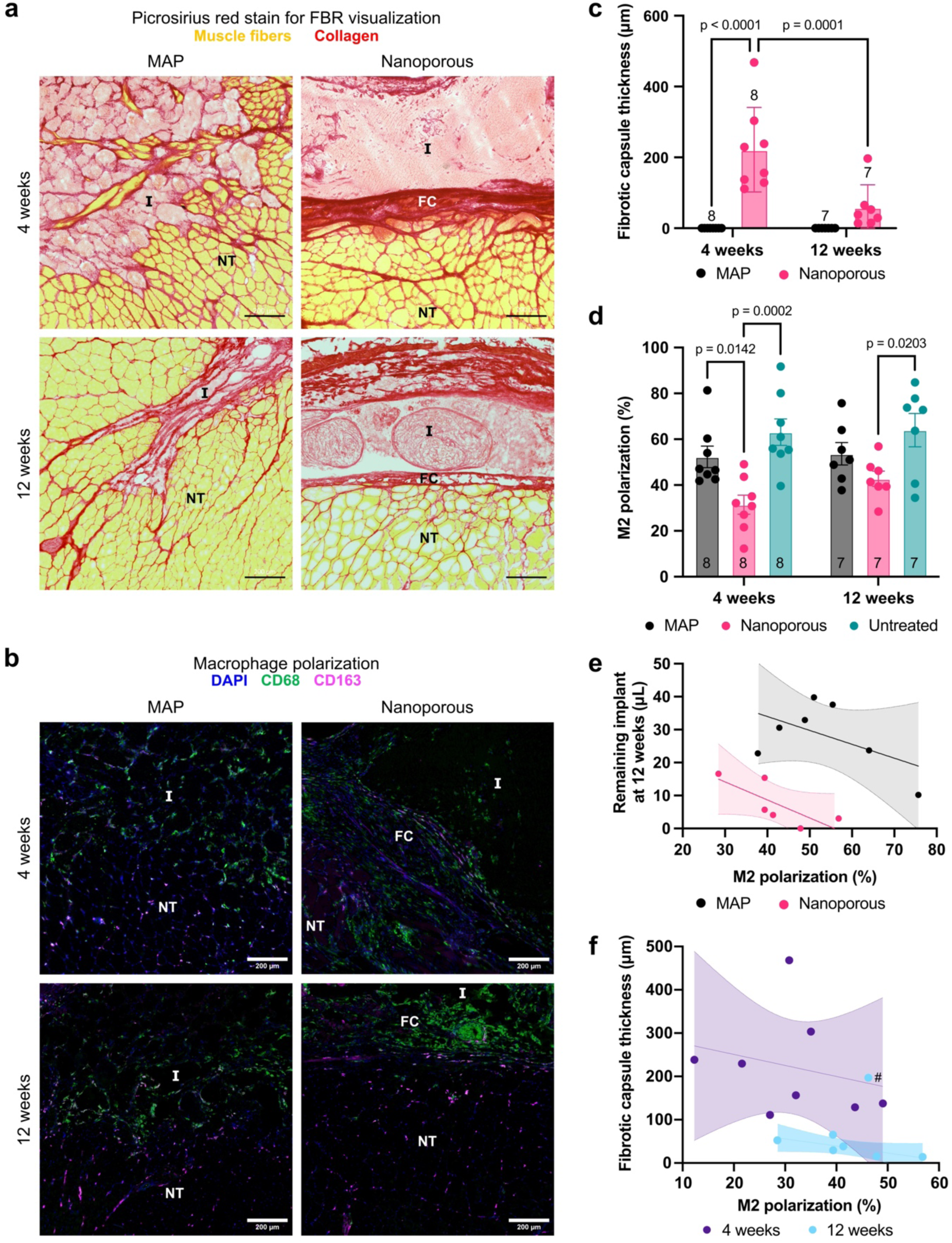
Local immune response to hydrogel implants. (a) Representative images of picrosirius red stain show fibrous capsule (indicated by arrow) formation around nanoporous implants, but not MAP implants. (b) Representative images of macrophage immunofluorescence stain, showing a high density of macrophages at the interface of nanoporous implants and uniform infiltration into MAP implants. (c) Quantification of fibrotic capsule thickness, showing a decrease in thickness from 4 weeks to 12 weeks in nanoporous implants, as well as no capsule formation for MAP implants. (d) Quantification of M2 macrophages (CD68^+^/CD163^+^), showing that nanoporous implants have significantly less M2 macrophages than MAP implants and untreated muscles. (e) M2 macrophage polarization is correlated with the volume of remaining implant at 12 weeks for nanoporous implants. (f) Fibrotic capsule thickness is correlated with M2 macrophage polarization at 12 weeks for nanoporous implants. Scale bars represent 200 μm. I = implant, FC = fibrotic capsule, NT = native tissue. # indicates an outlier identified by outlier analysis and excluded from statistical tests.

To assess the impact of scaffold porosity more directly on the local immune response, we quantified the polarization of macrophages using CD68 as a general macrophage marker and CD163 as a marker of the pro-regenerative phenotype (i.e., alternatively activated or M2 macrophages). Immunofluorescent staining (Fig. **4b**) showed that macrophage polarization is shifted further towards the M2 phenotype (CD68^+^/CD163^+^) in MAP scaffolds compared to nanoporous scaffolds, with a significant difference in the proportion of M2 macrophages between the hydrogel groups at 4 weeks (Fig. **4d**). Although statistical significance was lost by 12 weeks, MAP scaffolds still trended higher in M2 polarization at that timepoint. Nanoporous scaffolds also had significantly decreased M2 polarization compared to untreated animals at both timepoints, while MAP was statistically similar to the untreated group at both timepoints. Upon closer examination of the data, we found that the remaining volume of scaffold was negatively correlated to M2 polarization at 12 weeks for the nanoporous implants (p = 0.04, r = -0.76, r^2^ = 0.57) (Fig. **4e**). In addition, the thickness of the fibrotic capsule was also negatively correlated with the macrophage polarization of nanoporous implants at 12 weeks (p = 0.04, r = -0.75, r^2^ = 0.57) (Fig. **4f**). This would indicate that the local immune response plays an important role in the degradation behavior of the scaffolds, as well as the thickness of the fibrotic capsule.

### 2.5. MAP implants support angiogenesis and myogenesis

Blood vessels are vital support structures in skeletal muscle, providing oxygen and nutrients necessary for muscle contraction and removing waste products. After branching off arterioles, extensive capillary networks are formed in the extracellular matrix between muscle fibers, with most capillaries running parallel to the fibers. These microvascular networks are highly adaptable to muscle activity, such as exercise, remodeling to ensure energetic needs are met.^72, 73^ Accordingly, angiogenesis is an important component of regeneration after muscle injury, and it must be properly coordinated with myogenesis to enable the formation of functional muscle fibers.

To assess the impact of scaffold porosity on angiogenesis, we labeled blood vessels with CD31, a marker for endothelial cells (Fig. 5). We found a stark contrast in the organization of newly formed vasculature in the area of the defect, depending on the treatment. MAP scaffolds exhibited a moderate level of angiogenesis throughout the implant, with vessel diameters that were similar to that of capillaries in the surrounding tissue. This pattern was conserved across time points. Nanoporous implants, however, exhibited low angiogenesis within the hydrogel, with most new blood vessels forming at the perimeter. Larger blood vessels that were distinct from the usual capillary network were found in the fibrotic tissue surrounding the implant, and individual CD31^+^ cells were occasionally distinguishable within nanoporous implants. Interestingly, the surface of nanoporous implant had a high quantity of blood vessels that were derived from vasculature found in the fascia.

**Figure 5.**
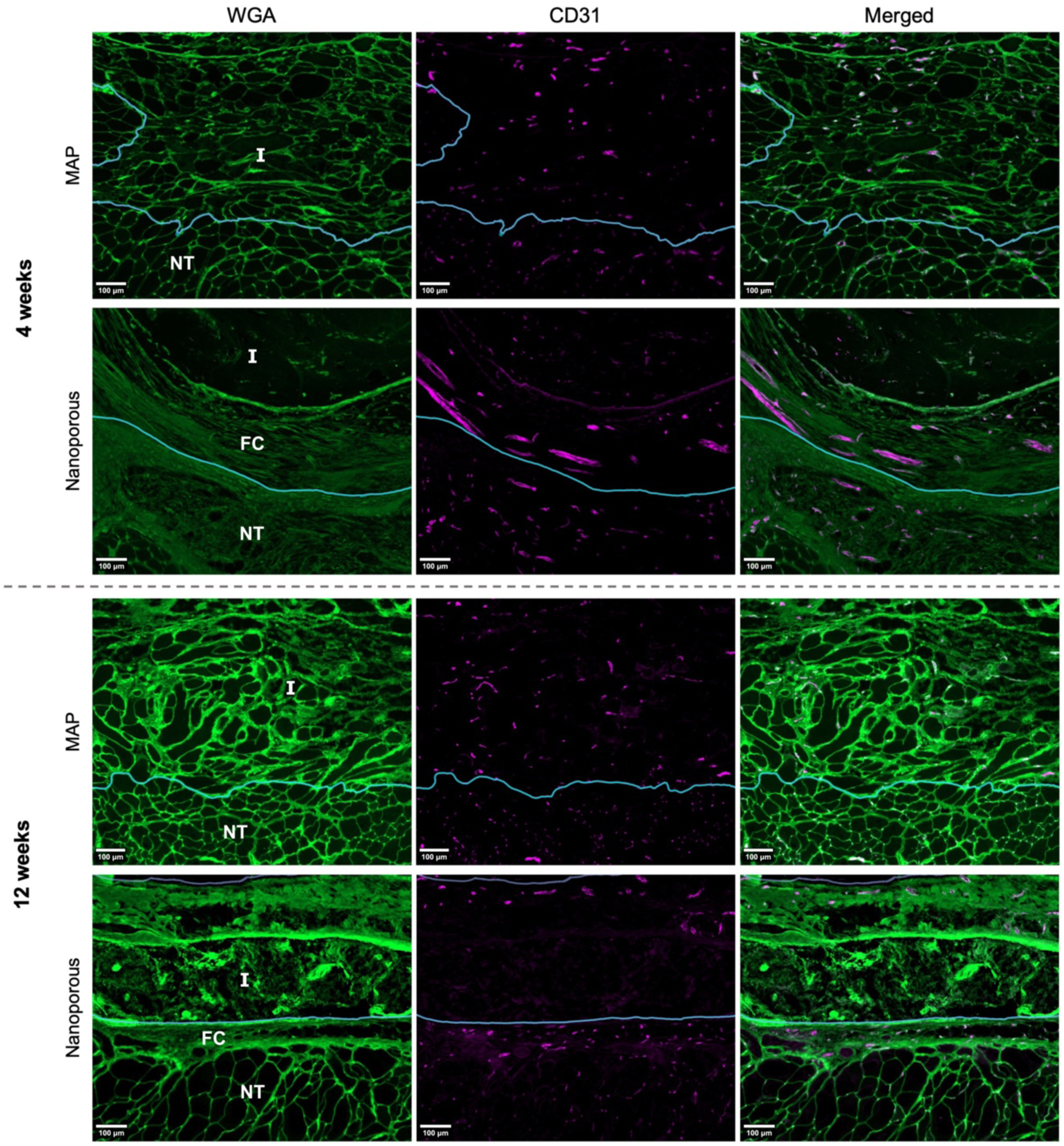
Angiogenesis in response to hydrogel implants. Representative images of WGA and immunofluorescent CD31 staining show formation of blood vessels throughout the MAP implants at both 4 weeks and 12 weeks. In contrast, angiogenesis in nanoporous implants occurred predominantly within the fibrotic tissue surrounding the implant and in the vicinity of the fascia, with little blood vessel penetration into the implant itself. In addition, large CD31^+^ structures were found in the fibrotic capsules of nanoporous implants, in contrast with the small capillaries that are characteristic of vasculature near the surface of the TA. Border of implant and native tissue demarcated by blue line. Areas below the line are native tissue. Scale bars represent 100 μm. I = implant, FC = fibrotic capsule, NT = native tissue

Next, we used MF20 antibody to label myosin heavy chain and identify myofibers within our implants (Fig. 6). We found myofiber formation within MAP implants as early as 4 weeks. While some myofibers within the implant seemed to be aligned with the uninjured muscle fibers, most did not exhibit proper organization. However, myofibers could be found at large distances from the implant edge and formed extensively at the perimeter of the scaffold. Unlike MAP implants, nanoporous implants rarely had distinguishable myofibers within the scaffold, with most new fibers forming within the fibrotic tissue surrounding the implant.

**Figure 6.**
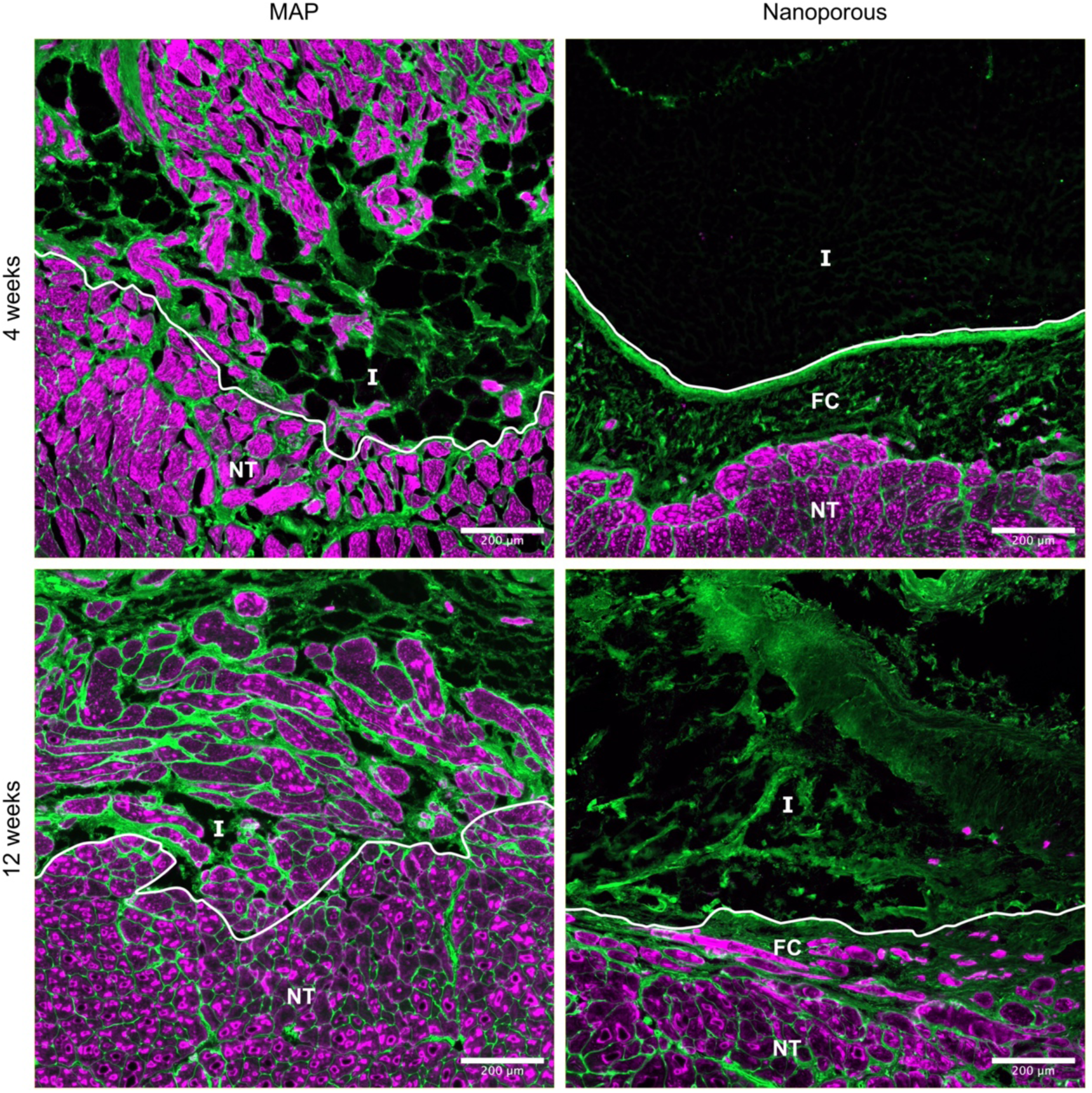
Myogenesis in response to hydrogel implants. Representative images of WGA (green) and immunofluorescent MF20 (red) staining show formation of muscle fibers within the MAP implants at both 4 weeks and 12 weeks. These new fibers form in the pore space between MAP microgels and are misaligned compared to pre-existing myofibers in the uninjured muscle. In contrast, very few new muscle fibers are found within nanoporous implants. Most newly formed myofibers in animals treated with a nanoporous scaffold are found in the fibrotic tissue surrounding the implant; therefore, the rate of muscle formation appears to be limited by the rate of degradation of the implant. Border of implant and native tissue demarcated by white line. Areas below the line are native tissue. Scale bars represent 200 μm. I = implant, FC = fibrotic capsule, NT = native tissue

### 2.6. MAP implants promote the formation of nerves and neuromuscular junctions

Neuromuscular units are severely disrupted by VML injuries, resulting in the denervation of muscle fibers. In a rodent model of VML in the tibialis anterior, the injury induced chronic axotomy of up to 69% of the motoneurons innervating this muscle, which worsened the functional deficit beyond what would be expected from the loss of muscle fibers alone.^74^ Muscle fibers create a microenvironment that promotes reinnervation at early stages post-denervation, but as time passes denervated muscle fibers and their NMJs become less receptive to reinnervation, resulting in muscle atrophy and increased fibrosis.^75^ Therefore, due to the time-sensitive nature of this regenerative process, biomaterial-based therapeutics for VML must create an environment that is conducive to reinnervation.

To evaluate the neuromuscular units in the defect area, NMJs were detected by staining for the nicotinic acetylcholine receptors of the postsynaptic endplate with α-bungarotoxin (α-BTX) and nerves were identified by immunostaining for the heavy chain of neurofilament protein (NF200) (Fig. **7a**). NMJ density was quantified for different regions of whole muscle cross-sections: implant region, interface region (<500 μm from implant/defect), and uninjured area (>1000 μm from implant/defect). The densities of the implant and interface regions were then normalized to that of the uninjured area in the same cross-section to account for the branching organization of neurovascular bundles in the TA, which can result in varying NMJ densities among cross-sections depending on the location of the cross-section along the TA’s longitudinal axis.^76^ NMJs were detected within the implants at significantly higher relative densities in MAP scaffolds compared with nanoporous scaffolds as early as 4 weeks, and this pattern was maintained at 12 weeks (Fig. **7b**), indicating that the newly formed myofibers found within MAP implants (Fig. **4a** and Figure 6) have been innervated. NMJs were found at statistically similar densities in the interface region across all groups at both timepoints (Fig. **7c**), indicating that the implants did not negatively impact NMJs compared to the untreated group.

**Figure 7.**
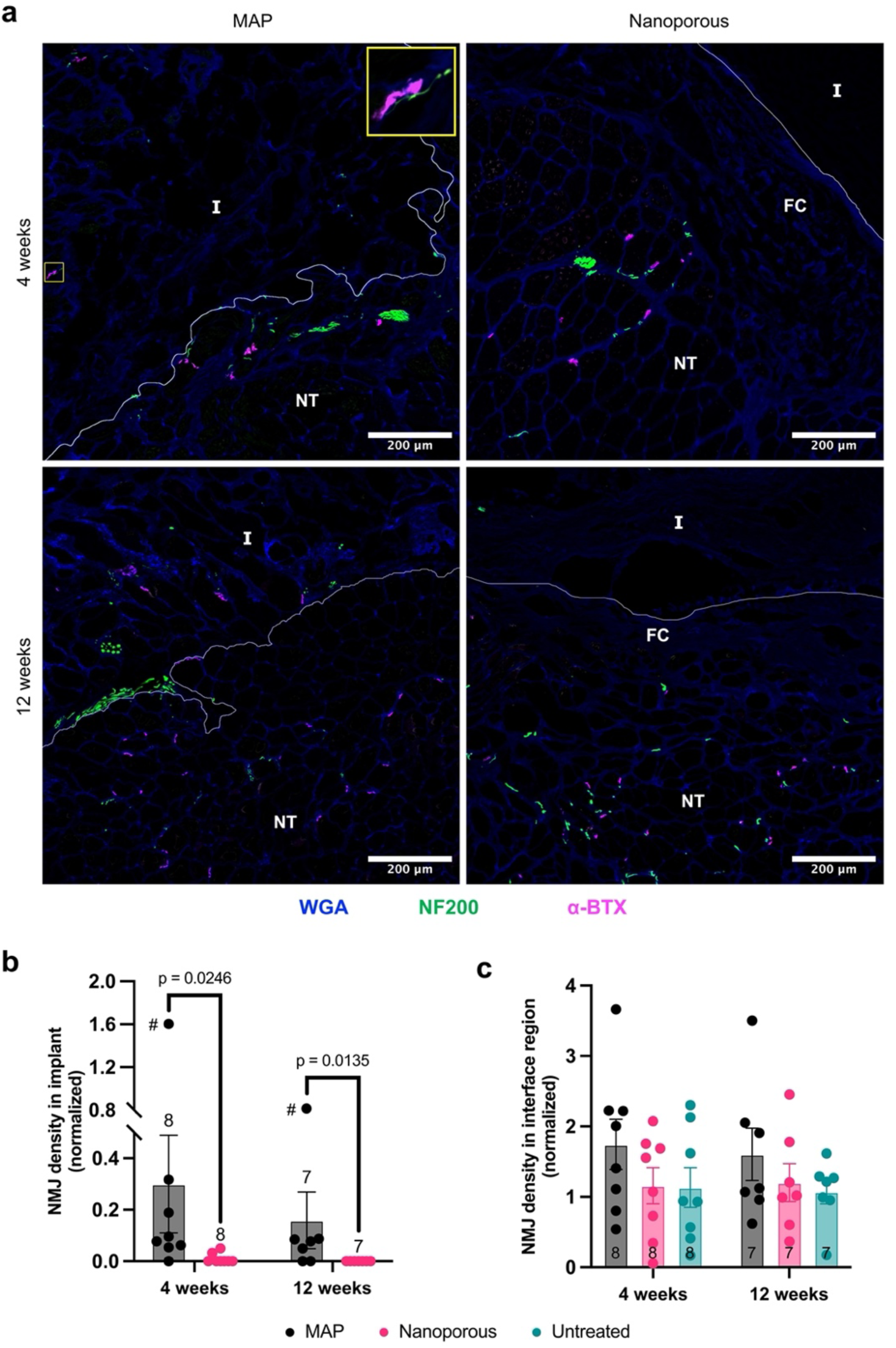
Neuromuscular junction formation in the defect area. (a) Representative images of the implant interface show NMJs within MAP implants and in the vicinity of both MAP and nanoporous implants. Inset shows example of an NMJ with its attached nerve within a MAP implant. (b) NMJ density within the implant relative to that of uninjured areas of the muscle is significantly higher for MAP implants compared to nanoporous implants at both timepoints. # indicates an outlier identified by outlier analysis and excluded from statistical tests. (c) NMJ density in the interface region (up to 500 μm from the edge of the implant for hydrogel treatments or from the surface of the muscle for untreated defects) relative to uninjured areas is statistically similar across groups at both timepoints. Each point in (b) and (c) represents the average value for an individual animal. Border of implant and native tissue demarcated by white line. Areas below the line are native tissue. Scale bars represent 200 μm. I = implant, FC = fibrotic capsule, NT = native tissue

Additional staining for important structures associated with the NMJ was performed to assess the stability of NMJs found within the defect. Representative images of thick tissue sections (50 μm) show that, along with the postsynaptic endplate (α-BTX), newly formed NMJs within defects treated with MAP had co-localized presynaptic endplates (SV2) and accompanying nerves (NF200) (Fig. **8a**). Schwann cells (S100), essential for the formation and maintenance of NMJs and myelinated nerves, were also found co-localized with the endplate and with associated nerves.

**Figure 8.**
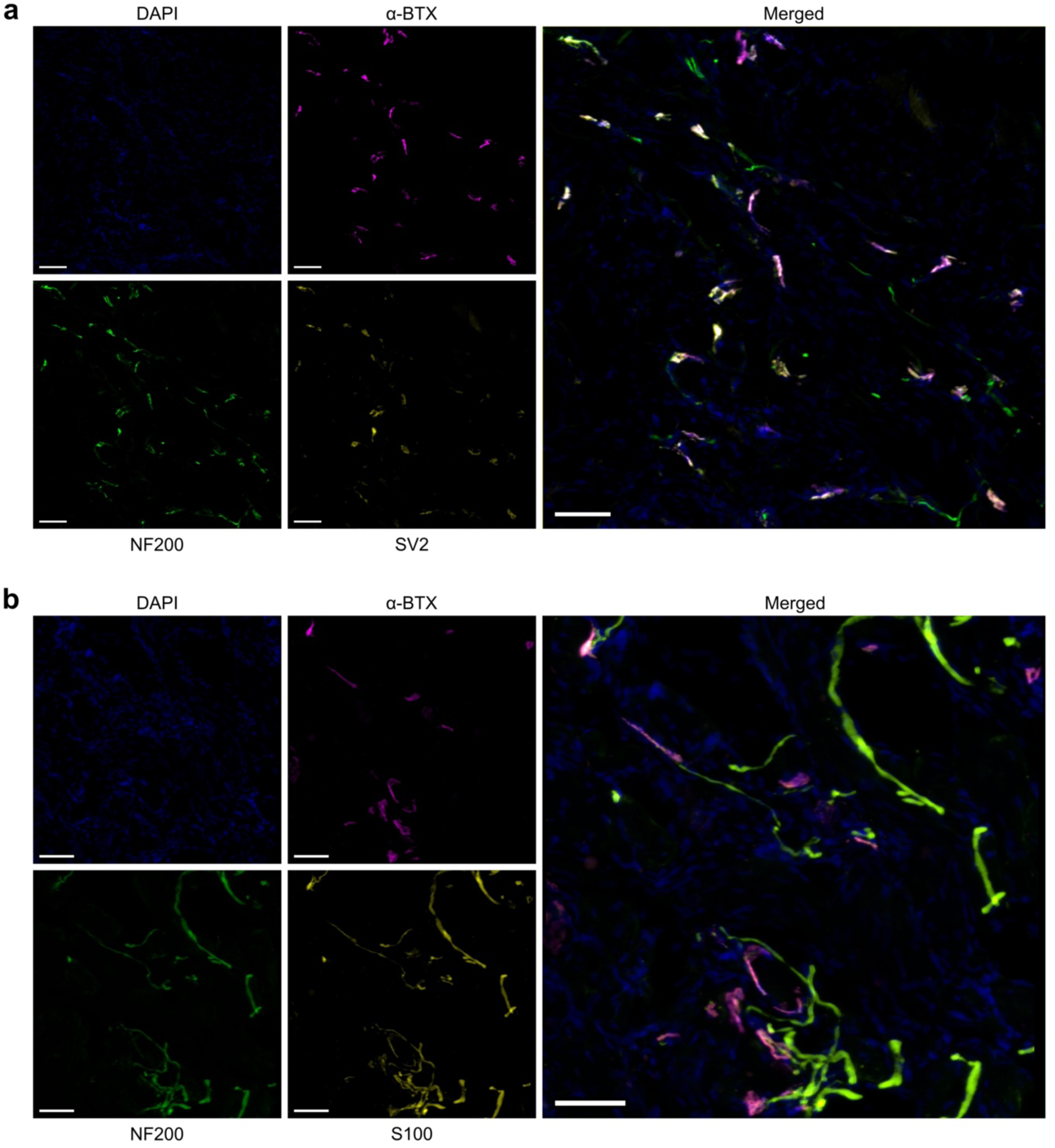
Neuromuscular junctions within defect treated with MAP scaffold exhibit appropriate structure. Representative images of NMJs within regenerated muscle tissue at 12 weeks post-treatment with MAP scaffold show that essential components for NMJ function are present. (a) In addition to the postsynaptic endplate found on the muscle fiber (α-BTX), NMJs in the regenerated tissue also possess nerves (NF200) and presynaptic endplates (SV2). (b) Schwann cells (S100) are present at the presynaptic terminal, where they support NMJ formation and function, and along the neural axon. Images shown are from consecutive sections. Scale bars represent 50 μm.

### 2.7 Functional recovery requires native-like muscle fiber alignment

A therapy for VML must result in meaningful functional recovery to be considered effective. Although MAP scaffolds promoted the formation of support structures and enabled extensive muscle fiber development in the defect, the TA did not recover its pre-injury function. The maximum torque achieved by each animal relative to its body weight was far below the baseline values (Fig. 2b) at all time points post-injury for all groups (Fig. **9a**). Following this trend, the ratio of functional recovery remained low for all groups (Fig. **9b**), with the only significant difference in either metric of function occurring between nanoporous and untreated groups.

**Figure 9.**
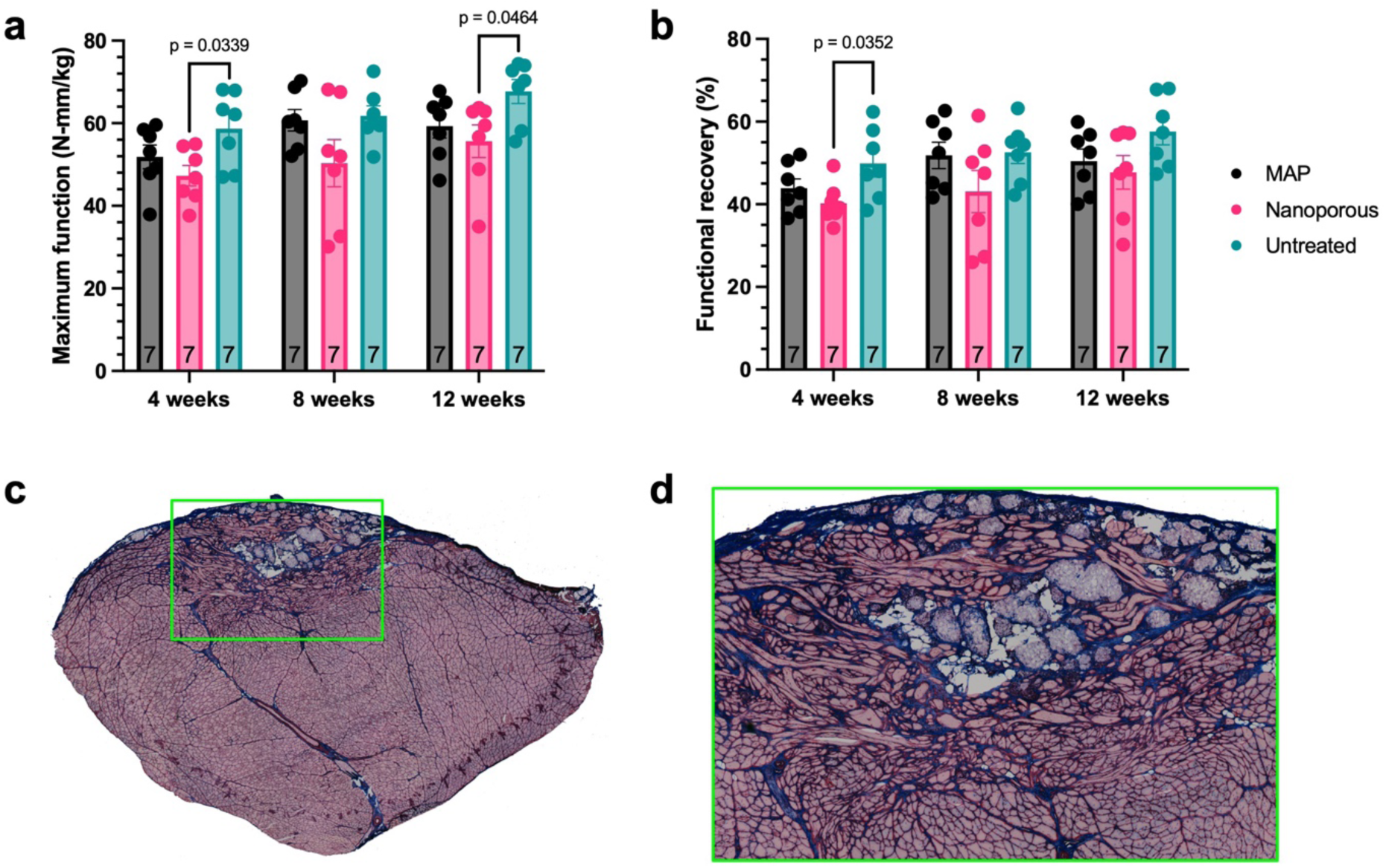
Functional recovery post-treatment. Measurements of torque production at 4 weeks, 8 weeks, and 12 weeks post-injury for the 12-week cohort show (a) diminished force production (normalized to body weight) at all timepoints compared to the baseline, resulting in (b) poor overall functional recovery for all treatment groups. (c) A representative image of late-stage regeneration in a muscle treated with MAP scaffold shows substantial myofiber formation in the defect. (d) An inset centered on the defect in (c) reveals considerable disorganization of myofibers, with only a small proportion aligned with the native muscle fibers.

This result emphasizes the need for tissue remodeling during the process of regeneration in skeletal muscle injuries to closely match the native tissue architecture and structure. *In vitro* studies have shown that muscle’s tightly bundled, anisotropic microarchitecture is an important factor for force production, to the extent that aligned myofibers can produce as much as 20 times greater force compared to unaligned myofibers.^77, 78^ Therefore, the force production of a muscle treated with MAP scaffold will not reflect the substantial muscle fiber formation within the defect (Fig**. 9c-d**) until the fibers become properly aligned.

## 3. Conclusion

Endogenous repair mechanisms in skeletal muscle are unable to overcome the challenges posed by volumetric muscle loss injuries, and no treatment available in the clinic has been able to effectively regenerate lost tissue and restore function to VML patients. The irreversible nature of the damage associated with VML, the maladaptive immune response to these injuries, and the chronic secondary denervation that follows further complicate the development of an effective therapeutic; it must be able to halt these deleterious processes as well as provide an environment conducive to the regeneration of necessary structures. MAP hydrogel is flowable and injectable before it is annealed to form a bulk scaffold, making it an ideal candidate for the treatment of irregularly shaped injuries of variable size, such as those characteristic of VML. In addition, microgels can be formed using any hydrogel formulation; therefore, MAP scaffolds are a flexible platform that can be used for modular biomaterial design in combination with other approaches.

In this study using MAP scaffolds to treat VML, we observed a significant effect on various processes modulated by the local immune response when the defect was treated with a microporous implant versus a nanoporous one. MAP scaffolds seemed to completely prevent the formation of a fibrotic capsule around the implant, indicating that the FBR has become negligible. The presence of fibers inside the MAP scaffold support this conclusion, showing that integration with the host tissue has been achieved. Macrophage polarization was shifted toward a more regenerative state in a microporous implant environment compared to a nanoporous environment, showing that microporosity is immunomodulatory in the context of VML. Notably, MAP scaffolds degraded significantly less than chemically identical nanoporous scaffolds, even though MAP scaffolds have less material to degrade in the same volume of hydrogel. Importantly, we report that MAP implants promote the formation of innervated muscle fibers within the implant as early as 4 weeks, as evidenced by the presence of NMJs on these fibers. However, due to the disorganization and misalignment of the newly formed fibers within the defect, we were unable to achieve meaningful functional recovery. We plan to conduct studies optimizing MAP properties and combining MAP scaffolds with physical rehabilitation to further promote remodeling of VML defects with an aim to also achieving improved functional recovery.

## 4. Methods

*Hydrogel formulation:* The precursor solution for microgel synthesis had a 3.0 wt% (w/v) polyethylene glycol (PEG) backbone and was prepared by mixing equal volumes of working solutions A and B. Working solution A consists of 5.64 mM 4-arm PEG-maleimide (10 kDa, Nippon Oil Foundry, Japan), 0.75 mM MethMal annealing macromer (synthesized as previously described^66^), and 2 mM RGD peptide (Ac-RGDSPGGC-NH2, Watson Bio). Working solution B consists of 10.78 mM enzymatically degradable crosslinker (Ac-GCGPQGIAGQDGCG-NH2, Watson Bio) and 10 μM biotin-PEG6-maleimide (TCI Chemicals). To synthesize microgels, working solution A was prepared using 10X PBS at pH 1.5, and working solution B was prepared using 1X PBS at pH 7.4. The two working solutions were then thoroughly mixed at a 1:1 ratio to create the precursor solution. To create the nanoporous implants, both working solutions were dissolved at twice their concentrations (solutions A2 and B2) in 1X PBS at pH 7.4 and aliquoted for use in surgery (40 μL for each aliquot). Aliquots (80 μL) of 1X PBS at pH 9.15 were also prepared. All solutions used for nanoporous implants were sterile filtered prior to aliquoting and frozen at -20 °C afterward.

*Mechanical matching:* Bulk hydrogel constructs were used to mechanically match the hydrogel formulation to skeletal muscle (12 kPa^68^). Small hydrogel disks (100 μL) were formed between two glass slides treated with SigmaCote that were spaced 2 mm apart. After gelation was completed, the hydrogel disks were swollen to equilibrium in 1X PBS overnight at 37 °C. Immediately before testing, the disks were removed from PBS and excess moisture was wicked away. The disks were then tested for compressive stiffness using an Instron Universal Testing System at a rate of 0.5 mm min^-1^. BlueHill software and the MATLAB SLM package were used to create stress-strain curves and calculate the Young’s modulus (Pa).

*Microgel size quantification:* A dilute solution of microgels (1:100) was prepared in 1X PBS with 1 μM fluorescently labeled streptavidin. After at least 15 min incubation at 37 °C, the solution was imaged in a 96-well plate (100 μL per well) using an ImageXpress Micro Confocal microscope (Molecular Devices) and particle diameters were quantified with an ImageXpress analysis module. Over 6000 particles were analyzed to determine average diameter and polydispersity index (PDI).

*Microgel synthesis:* Microgels were generated with a step-emulsification device (11.4 μm channel height) using a high-throughput method previously described^67^. A flow rate of 3.75 mL hr^-1^ was used for the precursor solution, while a flow rate of 7.5 mL hr^-1^ was used for the oil solution (2% Pico-Surf in NOVEC 7500). A solution of triethylamine (20 μL mL^-1^ precursor solution) dissolved in NOVEC 7500 was added to the collection tube to accelerate gelation. After gelation was finished, surfactant was removed from the microgels with successive washes in NOVEC 7500, 1X PBS, and hexanes as previously described^54^. Unreacted maleimides were quenched by incubating the microgels in 100 mM N-acetyl-L-cysteine (Acros Organics) overnight at 37 °C. Following three washes in 1X PBS, microgels were sterilized with three washes in 70% isopropyl alcohol (IPA) and stored in 70% IPA at 4 °C until further use.

*Preparation of MAP hydrogel for implantation:* Microgels were transitioned from 70% IPA to PBS with 5 successive washes in sterile 1X PBS. A sterile solution of 1 mM lithium phenyl-2,4,6-trimethylbenzoylphosphinate (LAP) in 1X PBS was prepared, and microgels were incubated in the photoinitiator solution at a 1:1 ratio (v/v) for at least 15 minutes. The microgels were then centrifuged at 18,000 g for 5 min and the supernatant was removed. Centrifugation and removal of supernatant was repeated until no supernatant could be visualized. The microgels were loaded into sterile 1 mL syringes and protected from light with aluminum foil.

*Animal care:* This study was conducted in compliance with the University of Virginia Animal Care and Use Committee guidelines and all procedures were approved under protocol 4045. We used a well characterized rodent model of VML in the tibialis anterior (TA)^71^. Twelve week-old female Lewis rats were purchased from Charles River and used for all studies. Rats were pair-housed in a vivarium accredited by the American Association for the Accreditation of Laboratory Animal Care and provided with enrichment, food, and water *ad libitum*.

*In vivo functional testing:* We evaluated TA function before defect creation to establish a baseline and at 4, 8, and 12 weeks after surgery as previously described^79^. Briefly, rats were anesthetized (1.5-2.5% isoflurane) and placed in a supine position on a heated platform with their knee stabilized at a 90° angle and their foot secured to the footplate of the Aurora Scientific 305-LR-FP servomotor. Two percutaneous needle electrodes were superficially inserted into the skin of the anterior compartment of the lower leg along the peroneal nerve. Electrical stimulation was applied in a controlled manner using the Aurora Scientific Stimulator Model 701C and Dynamic Muscle Control software at a range of frequencies (1-200 Hz). Functional testing was performed on the left hindlimb only.

*VML defect creation and treatment:* Surgical creation was performed as previously reported. Briefly, rats were anesthetized (2-2.5% isoflurane), administered analgesia (Ethiqa XR, 0.65 mg kg^-1^), and the left hindlimb was aseptically prepared. A longitudinal incision was made on the lateral aspect of the lower leg, and the skin was separated from the fascia. Another longitudinal incision was made along the lateral aspect the fascia over the TA. Blunt dissection was used to gently separate the fascia from the TA and expose the anterior crural muscles. The extensor digitorum longus (EDL) and extensor hallucis longus (EHL) were isolated and removed. The boundaries of the middle third of the TA (longitudinal axis, approximately 1 cm in length) were marked with a sterile surgical marker. Having previously calculated the theoretical weight of the TA (0.0017 × body weight), approximately 20% of the TA was removed, leaving 2-3 mm margins on the lateral and medial sides of the muscle and avoiding damage to the distal tendon. For animals treated with MAP scaffolds, 100 μL of microgels were injected into the defect and irradiated with 365 nm LED light (ThorLabs, 440 mA) for 5-10 s, until the scaffold was annealed. For animals treated with nanoporous implants, one aliquot each of working solution A2, working solution B2, and pH 9.15 1X PBS were thawed and their entire volumes were thoroughly mixed immediately before applying 100 μL of the mixture to the defect. Gelation of the nanoporous gel was verified before proceeding. The fascia was closed with interrupted sutures and 6-0 vicryl. The skin was closed with interrupted sutures and 5-0 prolene, and a thin layer of skin glue was applied over the skin sutures.

*MRI quantification of implant:* Animals were scanned at 4 weeks and 12 weeks post-surgery with a T2-mapping sequence using a small animal MRI machine (7 Tesla Bruker/Siemens ClinScan). Rats were anesthetized (1.5-2.5% isoflurane) and images were acquired from knee to ankle with slice thickness of 0.7 mm. No contrast agent was used. The images were analyzed on OSiriX to calculate the total volume of hydrogel implant remaining in the defect. One animal from the nanoporous group in the 12 week cohort was not imaged due to scheduling constraints at the core facility.

*Tissue collection:* At the endpoints, animals were anesthetized by CO2 inhalation. TA muscles were isolated, cut in half through the middle of defect (perpendicular to the longitudinal axis), and immediately snap-frozen in liquid nitrogen-cooled isopentane (- 150 °C). Samples were then embedded in OCT. Cryosections of 10 μm thickness were obtained, with consecutive sections on the same slide obtained from locations at least 200 μm apart on the block. Cryosections of 50 μm thickness were obtained for additional NMJ characterization.

*Picrosirius red staining*: Slides were rehydrated through a graded series of EtOH (100%, 95%, 70%; 3 min each) to deionized water and fixed in 4% PFA for 15 min before staining with picrosirius red according to the manufacturer’s instructions (Abcam, cat#: ab150681). *Immunostaining:* Slides were allowed to warm to room temperature before staining. For Pax7 and macrophage stains (CD68 and CD163), we fixed the slides for 10 min in 4% PFA and then performed heat induced antigen retrieval. We submerged the slides in a citrate-based unmasking solution (Vector Laboratories, cat#: H-3300) and processed them in an Instant Pot pressure cooker for 20 minutes on the low pressure setting with natural release. The slides were then left to cool down to room temperature for 1 hr. Slides were rinsed with PBS and sections were circled with a hydrophobic pen (Vector Laboratories, H-4000). Slides were washed with 10 mM glycine in PBS for 10 min, then washed twice with 0.5% Tween 20 in PBS. They were incubated for 30 min in blocking buffer^80^ (2% goat serum, 50 mM glycine, 0.05% Tween 20, 0.1% Triton X-100, and 0.1% BSA in PBS) at room temperature, followed by overnight incubation with the primary antibody at 4 °C. Primary antibodies were diluted in antigen signal enhancement solution^80^ (10 mM glycine, 0.05% Tween 20, 0.1% Triton X-100, 0.1% hydrogen peroxide). Slides were washed in 0.5% Tween 20 for 5 min, then washed twice in PBS for 5 min. Secondary antibodies were diluted 1:1000 in 0.1% Tween 20, and slides were incubated overnight at 4 °C. Slides were washed four times in PBS for 5 min, incubated with DAPI (Thermo Scientific, cat#: 62248; 1:1000 in PBS) and wheat germ agglutinin (WGA) (Invitrogen, cat#: W32464; 5 μg mL^-1^) for 30 min, and washed three times in PBS for 5 min. Slides were mounted with Prolong Gold (Invitrogen, P36930). For all other antibodies, slides were fixed in cold 100% MeOH for 10 min at 20 °C, followed by three PBS rinses (5 min each). Sections were circled with a hydrophobic pen, then slides were permeabilized in PBS-T (0.1% Triton X-100 in PBS) for 10 min. Slides were incubated in Sea Block (Thermo Scientific, cat#: 37527) for 30 min, then incubated with primary antibodies overnight at 4 °C. Primary antibodies were diluted in 5% Sea Block in PBS-T. Slides were washed in PBS-T for 5 min, then washed twice in PBS for 5 min. Secondary antibodies were diluted 1:1000 in PBS-T, and slides were incubated overnight at 4 °C. Slides were washed four times in PBS for 5 min, incubated with DAPI (1:1000 in PBS) and WGA (5 μg mL^-1^) for 30 min, and washed three times in PBS for 5 min. Slides were mounted with Prolong Gold. For slides stained with NF200, an endogenous biotin blocking kit (Invitrogen, cat#: R37628) was used according to manufacturer instructions prior to incubation with primary antibodies. Biotinylated α-Bungarotoxin (BTX) (Invitrogen, cat#: B1196) was included in the primary antibody diluent (1 μg mL^-1^) to label neuromuscular junctions (NMJs), and fluorescently labeled streptavidin (Invitrogen, cat#: 84547) was included in the secondary antibody diluent (1 μg mL^-1^). Slides were imaged with a 20x objective on an ImageXpress Micro Confocal microscope (Molecular Devices) or on an Axioscan 7 Slide Scanner (ZEISS). Slides with 50 μm cryosections were imaged with a Leica DMi8 inverted microscope using a 40x oil immersion objective.

**Table 1.**
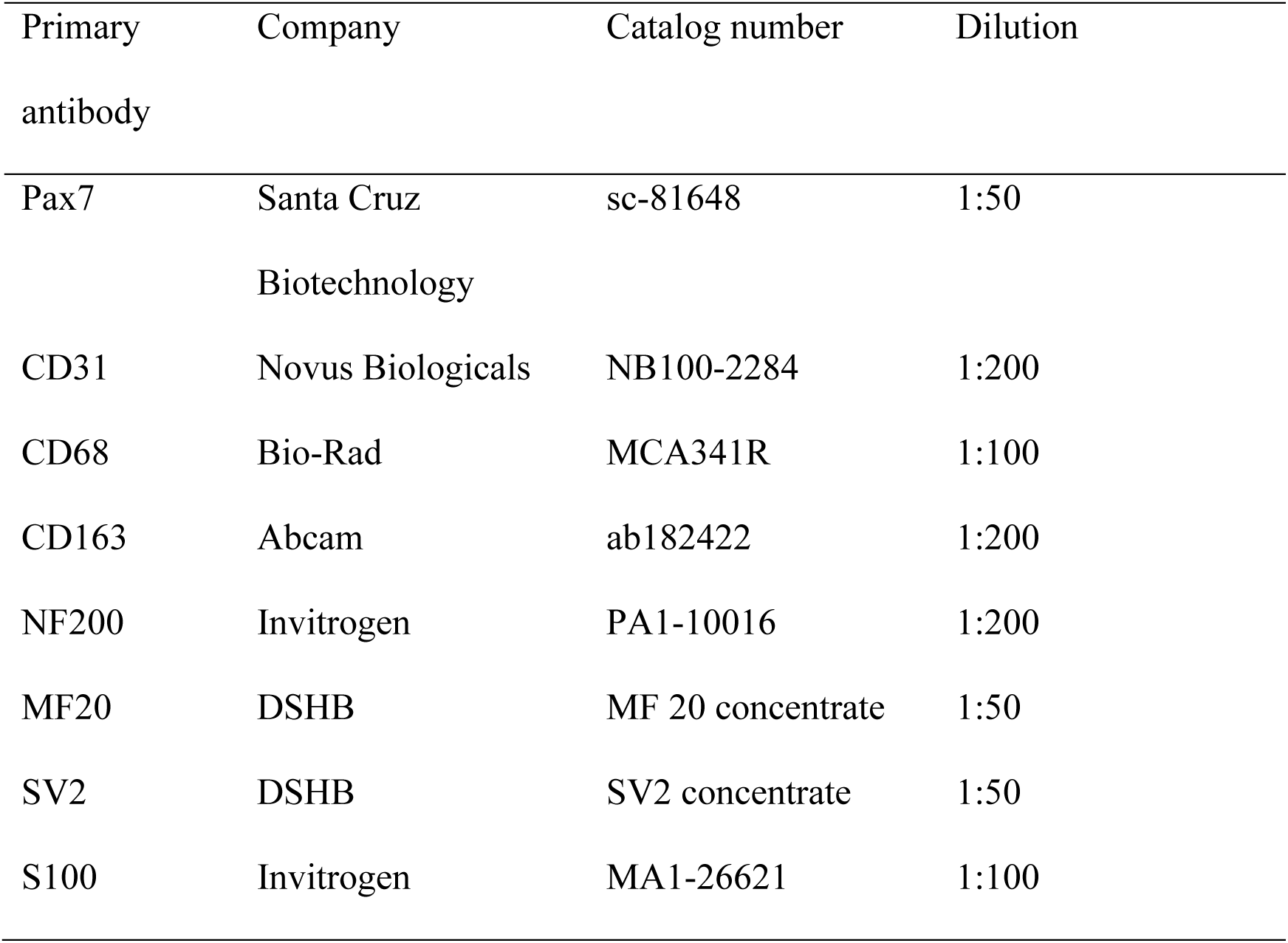
((Primary antibodies used for immunohistochemistry))

**Table 2.**
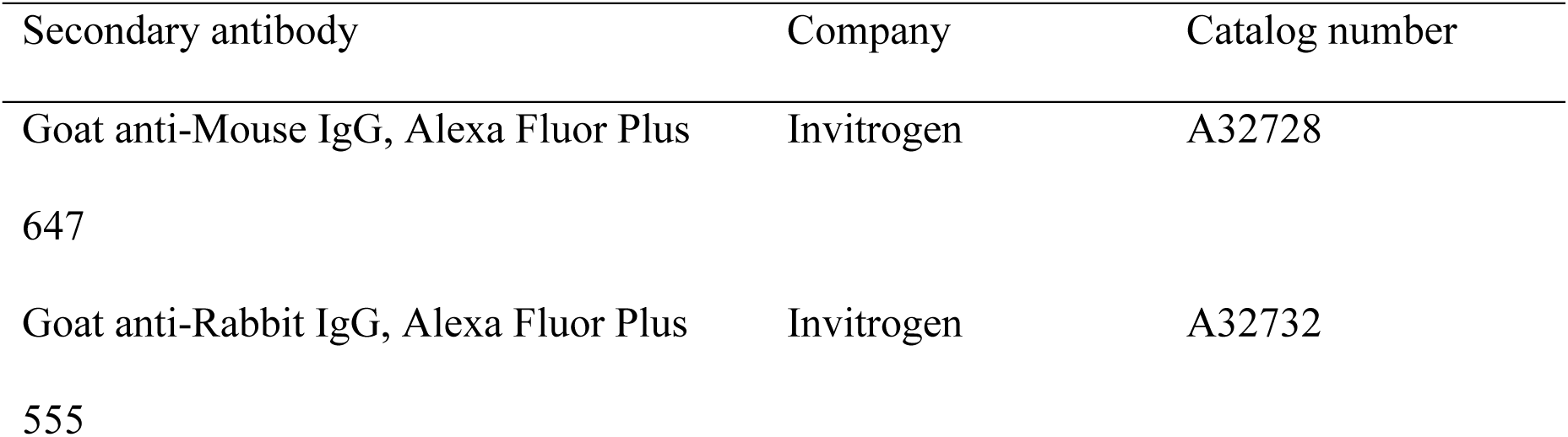
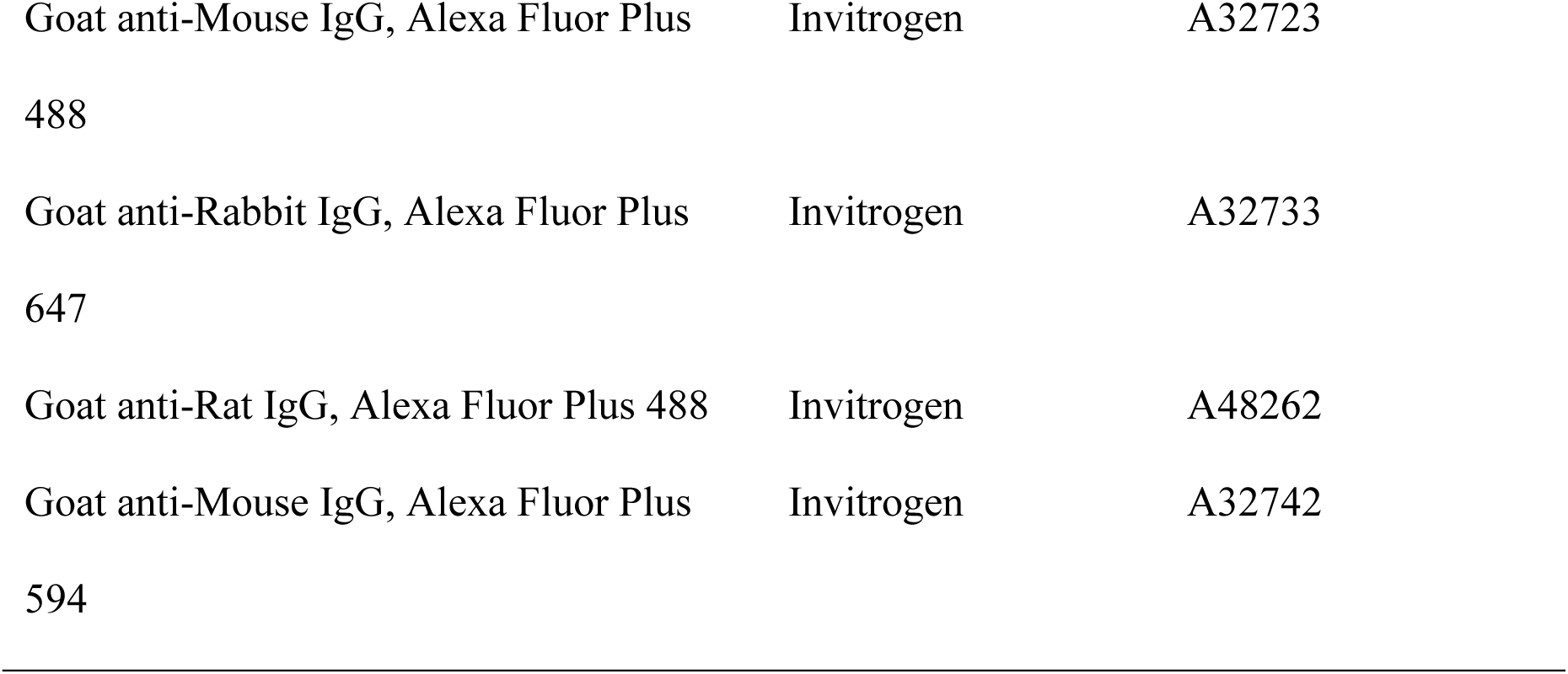
((Secondary antibodies used for immunohistochemistry))

| Secondary antibody | Company | Catalog number |
| --- | --- | --- |
| Goat anti-Mouse IgG, Alexa Fluor Plus<br>647 | Invitrogen | A32728 |
| Goat anti-Rabbit IgG, Alexa Fluor Plus<br>555 | Invitrogen | A32732 |
| Goat anti-Mouse IgG, Alexa Fluor Plus<br>488 | Invitrogen | A32723 |
| Goat anti-Rabbit IgG, Alexa Fluor Plus<br>647 | Invitrogen | A32733 |
| Goat anti-Rat IgG, Alexa Fluor Plus 488 | Invitrogen | A48262 |
| Goat anti-Mouse IgG, Alexa Fluor Plus<br>594 | Invitrogen | A32742 |

*Quantification of immune cells:* Slides were imaged with a 20x objective on an ImageXpress Micro Confocal microscope (Molecular Devices), and stitched images for whole sections were exported as TIF files. The MATLAB Image Processing Toolbox and a custom code was used to isolate square 0.5 mm^2^ regions of interest from whole cross-section images. At least three sections were analyzed per animal, and three random regions of interest within the area of the defect were analyzed per section. Quantification of nuclei, CD68^+^ cells and CD68^+^/CD163^+^ cells was performed using CellProfiler.^81^

*Quantification of fibrotic capsules:* Slides stained with picrosirius red were imaged with on a Leica DMi8 microscope, and stitched images for whole sections were exported as TIF files. At least three sections were analyzed per animal. Masks of fibrotic capsules were created and quantification of the thickness of fibrotic capsules was performed using the MATLAB Image Processing Toolbox.

*Quantification of nerves and NMJs:* Slides stained with NF200 and BTX were imaged with a 20x objective on an AxioScan 7 Slide Scanner (ZEISS), and stitched images for whole sections were exported as CZI files. Image analysis was performed on QuPath^82^. Masks were created for the region of the implant/defect, and another mask was extended 500 μm into the muscle for the interface region. At least three tissue sections were analyzed per animal. The watershed cell detection module was used to identify nerves (NF200^+^ areas) and neuromuscular junctions (BTX^+^ areas) in the whole section, and the masks were used to quantify features by location. Data exported from QuPath was analyzed with RStudio. The density of NMJs was calculated for each region, then normalized to the density of the uninjured region of each tissue section.

*Characterization of NMJs*: Slides stained with BTX, NF200, S100, SV2, and DAPI were imaged at 40x with a Leica DMi8 microscope to generate z-stacks (1 μm slices) of regions of interest. Average intensity projections were created for visualization.

*Quantification of blood vessels:* Slides stained with CD31 were imaged with a 20x objective on an AxioScan 7 Slide Scanner (ZEISS), and stitched images for whole sections were exported as CZI files. Image analysis was performed on QuPath^82^. Masks were created for the region of the implant/defect, and another mask was extended 500 μm into the muscle for the interface region. At least three tissue sections were analyzed per animal. A machine learning approach was used to train a pixel classifier for the identification of blood vessels (CD31^+^ structures) in the whole section. A training image “collage” composed of least three regions of interest per animal was generated and annotated manually, followed by the creation of a Random Trees model. The image features used in the model were Gaussian filter, Laplacian of Gaussian, Weighted Deviation, and Gradient Magnitude. Only data from the channels corresponding to WGA and CD31 were included, using full resolution (0.35 μm/px) and 0.5 Gaussian scale. Data exported from QuPath was parsed and summarized with RStudio.

*Quantification of satellite cells:* Slides stained with Pax7 were imaged with a 20x objective on an AxioScan 7 Slide Scanner (ZEISS), and stitched images for whole sections were exported as CZI files. At least three sections were analyzed per animal, and 5-8 random regions of interest within the area of the defect were analyzed per section. QuPath^82^ was used to quantify Pax7^+^ nuclei using the cell detection module. Data exported from QuPath was parsed and summarized with RStudio.

*Quantification of muscle fibers:* Slides stained with MF20 were imaged with a 20x objective on an AxioScan 7 Slide Scanner (ZEISS), and stitched images for whole sections were exported as CZI files. Image analysis was performed on QuPath^82^. Masks were created for the region of the implant/defect, and another mask was extended 500 μm into the muscle for the interface region. At least three tissue sections were analyzed per animal. The threshold pixel classifier was used to identify muscle fibers (MF20^+^) in the whole section, and the masks were used to quantify the percentage of positively labeled pixels by location. Data exported from QuPath was parsed and summarized with RStudio.

*Statistics:* Statistical analyses were performed in GraphPad Prism 10. All values are expressed as the mean ± standard deviation. The ROUT method was used for outlier analysis.

## Supporting Information

### Code Availability

The underlying code for this study is not publicly available but may be made available to qualified researchers upon reasonable request.

### Data Availability

Data supporting the findings of this study are available from the corresponding author upon reasonable request.

## Acknowledgements

This work was funded by the Center for Advanced Biomanufacturing at the University of Virginia.

